# Aerial course stabilization is impaired in motion-blind flies

**DOI:** 10.1101/2020.12.16.423059

**Authors:** Maria-Bianca Leonte, Aljoscha Leonhardt, Alexander Borst, Alex S. Mauss

## Abstract

Visual motion detection is among the best understood neuronal computations. One assumed behavioural role is to detect self-motion and to counteract involuntary course deviations, extensively investigated in tethered walking or flying flies. In free flight, however, any deviation from a straight course is signalled by both the visual system as well as by proprioceptive mechanoreceptors called ‘halteres’, which are the equivalent of the vestibular system in vertebrates. Therefore, it is yet unclear to what extent motion vision contributes to course control, or whether straight flight is completely controlled by proprioceptive feedback from the halteres. To answer these questions, we genetically rendered flies motion-blind by blocking their primary motion-sensitive neurons and quantified their free-flight performance. We found that such flies have difficulties maintaining a straight flight trajectory, much like control flies in the dark. By unilateral wing clipping, we generated an asymmetry in propulsory force and tested the ability of flies to compensate for this perturbation. While wild-type flies showed a remarkable level of compensation, motion-blind animals exhibited pronounced circling behaviour. Our results therefore unequivocally demonstrate that motion vision is necessary to fly straight under realistic conditions.

## Introduction

The coordination of muscle activity underlying locomotion is subject to internal and external perturbations inevitably leading to error accumulation over time. To control movement trajectories therefore requires continuous sensory feedback for adjustments (Taylor & Krapp, 2007; Rossignol et al., 2006; Dickinson, 2014; Tuthill & Azim, 2018). Vision is particularly well suited to keep the animal on track since any self-motion evokes characteristic image movements across the eye, termed optic flow (Gibson, 1950; Koenderink & van Doorn, 1987).

To investigate the influence of optic flow signals on course control, locomotor behaviours of various animals have been probed with a wide range of visual motion stimuli (Goetz, 1975; Collett & Land, 1975; Rock & Smith, 1986; Warren et al., 2001; Mronz & Lehmann, 2008). However, drawing conclusions from such experiments is non-trivial because the sensation-action feedback cycle underlying natural behaviour is usually broken to some extent. Simply depriving animals of their visual sense is not satisfactory, because positional cues, which may control heading independently of visual motion, are also removed. Although the neuronal basis of visual motion detection is understood in detail (Mauss et al., 2017; Wei, 2018), its use in course control is still unclear.

Flight is particularly interesting in the context of course stabilization since animals must control their locomotor trajectories fast and in three spatial dimensions (Egelhaaf, 2013). Flies are among the most agile flying animals requiring minute coordinated changes in all aspects of wing motion to control course (Muijres et al., 2014). A great demand on stabilizing sensory feedback is therefore expected (Egelhaaf, 2013; Dickinson & Muijres, 2016). Recently, it became possible to render flies motion-blind by selective genetic manipulation of the first stage of visual motion detection (Maisak et al., 2013; Bahl et al., 2013). Importantly, the visual position system (Bahl et al., 2013) and all other sensory modalities remain functional. Here, we take advantage of this experimental system to verify the long-held notion that visual motion feedback is required for course stabilization.

## Results and Discussion

### Probing free-flight

We released flies in a transparent enclosure (Collett & Land, 1975; Straw et al., 2011; Tammero & Dickinson, 2002) (Figure 1A) and tracked their positions in 3D at 100 frames per second using a calibrated camera system. We calculated features frame-by-frame, such as flight velocity and turning angle. Static or moving visual patterns were displayed at 120 Hz on four monitors surrounding the enclosure. We used wild-caught flies for an initial characterization. In agreement with previous accounts, recorded trajectories consisted of straight segments interspersed by sharp turns, so-called body saccades (Collett & Land, 1975; Mronz & Lehmann, 2008; Schilstra & van Hateren, 1999; Tammero & Dickinson, 2002; Egelhaaf et al., 2012) (Figure 1C, D). We detected saccades on the basis of turning velocity. Another characteristic feature of a saccade is a brief drop in flight velocity (Mronz & Lehmann, 2008; Schilstra & van Hateren, 1999; Tammero & Dickinson, 2002) (Figure 1D).

**Figure 1.**
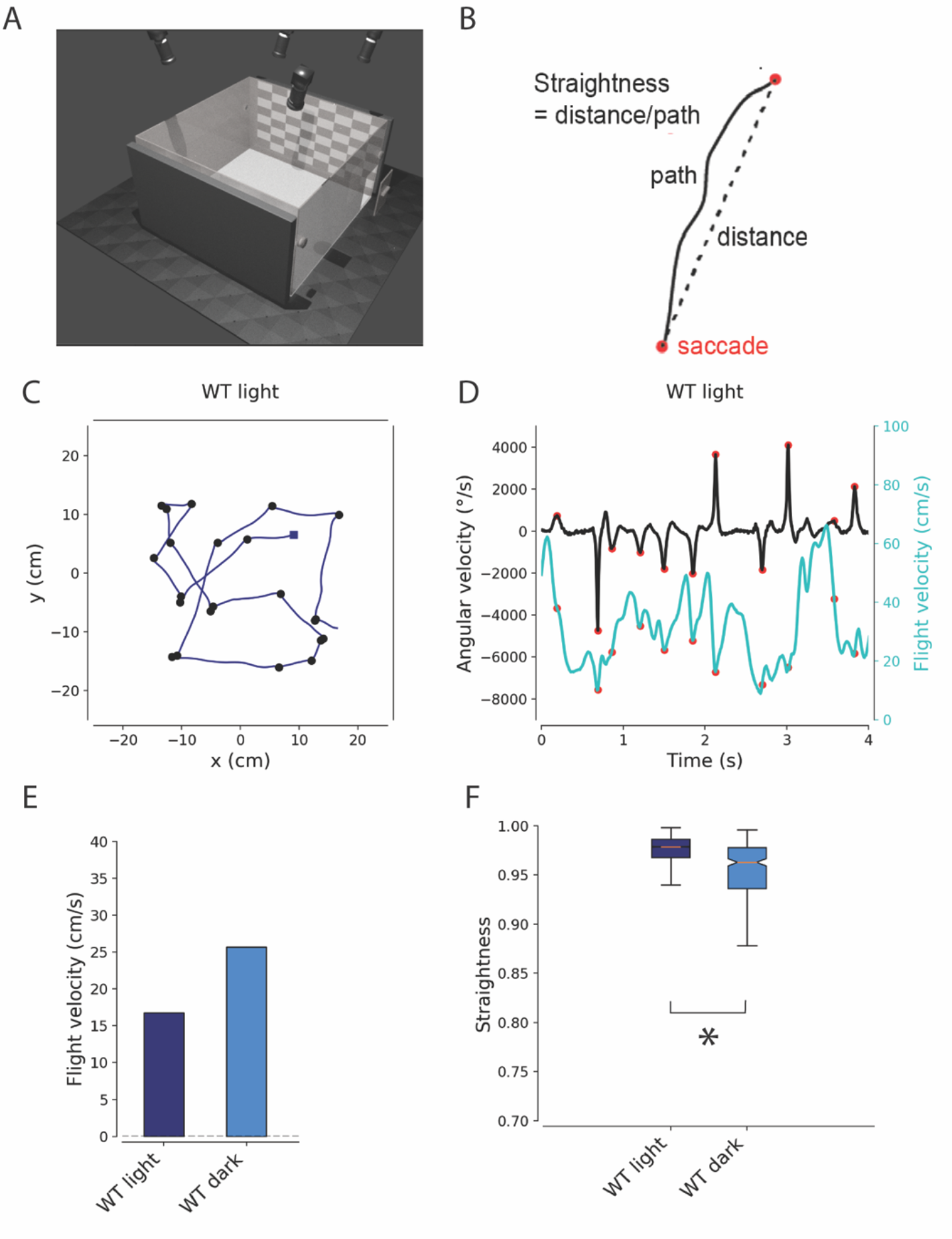
Free-flight behavior of wildtype in the light and dark. (**A**) Schematic representation of the free-flight arena: A plexiglass box (50×50×30) is surrounded by 4 monitors to display the visual stimuli. The arena is illuminated from underneath by infrared lights. Five cameras fitted with infrared filters record the flies from different angles. (**B**) The straightness index is calculated by dividing the distance between two consecutive saccades by the covered path length. A perfectly straight segment would yield a value of 1.0, while for a trajectory of the shape of a semicircle, the index would be 0.63.(**C**) Example of a single flight trajectory of a wild type fly projected in the horizontal plane. (**D**) Horizontal angular velocity and flight velocity values over time. Saccades (red) were identified based on angular velocity (peak values above 300 0/s). (**E**) Average flight velocity (s.e.m ≤ 0.045) of flies in the light and in the dark (n=1988 trajectories light and n=310 trajectories dark). (F) Straightness of long intersaccadic segments (250 – 2000 ms) of wild type flies flying in a lit (dark blue) or in a dark (light blue) arena. Significant difference is based on a Kolgomorov-Smirnov test (p<0.0001, n=4609 segments light and n= 302 segments dark)

### Flight with and without vision

Appropriate detection of various self-evoked optic flow components by neural circuits is instrumental in monocular depth perception (Srinivasan, 2011; Ravi et al., 2019; Egelhaaf et al., 2012) as well as estimating and regulating locomotor speed (Baird et al., 2005; Srinivasan et al., 1996; Pfeffer & Wittlinger, 2016). Since long, self-induced image motion is also thought to provide sensory cues to detect and counteract involuntary course deviations (Gibson, 1950; Goetz, 1975; Collett & Land, 1975; Egelhaaf, 2013).

We first asked how missing visual feedback affected flight structure by comparing trajectories of the same wild-type strain under two conditions: with visual patterns surrounding the arena (i.e. with intact visual feedback) and in darkness (i.e. without any visual feedback). Trajectories obtained in the dark were usually shorter in duration. In addition, the average flight velocity in darkness was increased (Figure 1E), in line with the idea that flight velocity is reflexively modulated by the received optic flow (Mauss & Borst, 2020; Baird et al., 2005; Srinivasan et al., 1996). All saccade metrics, however, were highly similar (Figure S1). This finding further supports the notion that, although visual signals can trigger saccades, the execution of a saccade is not under visual control (Bender & Dickinson, 2006; Tammero & Dickinson, 2002; Karmeier et al., 2006).

Closer inspection of flight trajectories revealed that vision is more important to maintain a stable bearing during inter-saccadic flight. To quantify this, we calculated a straightness index by dividing the distance between two consecutive saccades by the covered path length (Figure 1B). For each experimental condition, we further divided inter-saccadic segments into two groups: short (50 – 250 ms) and long (250 – 2000 ms). Comparing straightness indices between bright and dark condition revealed a significant reduction for long segments recorded in the dark (Figure 1F). We further obtained data from a completely blind fly strain, NorpA, with a mutation in the essential phototransduction enzyme Phospholipase C (Hotta & Benzer, 1970). NorpA flies flew even faster and less straight than wild-type flies in the dark (Figure S2).

### Free-flight behaviour of motion-blind flies

From the results above, we can conclude that vision is important for course stabilization. However, the question remains whether the stabilizing influence is exerted by visual motion signals. Course stabilization can also be achieved by keeping conspicuous visual features at a constant position on the retina (Bahl et al., 2013; Bar et al., 2015).

In the fly brain, optic flow signals are computed in several, hierarchical steps. First, the direction of visual motion is computed by T4 and T5 cells locally for each point in visual space (Maisak et al., 2013). Next, higher-order neurons collect the signals from sub-samples of appropriately aligned T4 and T5 cells to detect the presence of specific flow field components such as rotation, translation (Karmeier et al., 2006; Krapp & Hengstenberg, 1996) or expansion (Klapoetke et al., 2017). Any of such self-induced flow components may influence movement trajectories (Collett & Land, 1975; Rock & Smith, 1986; Warren et al., 2001; Mronz & Lehmann, 2008).

To test for the involvement of motion vision, we rendered flies motion-blind by cell-specific expression of tetanus toxin in the primary motion-sensing neurons (‘T4T5>TNT’). We confirmed the absence of motion-vision in these flies in two different ways. First, we found that the optomotor turning response of tethered walking flies was completely abolished (Figure S4). Second, we measured turning of freely flying flies in response to horizontal pattern rotation around the arena. In contrast to controls showing the expected following reaction (Mronz & Lehmann, 2008), responses of T4T5>TNT flies to moving patterns (clockwise and counter clockwise) did not reveal any turning bias (Figure 2).

**Figure 2.**
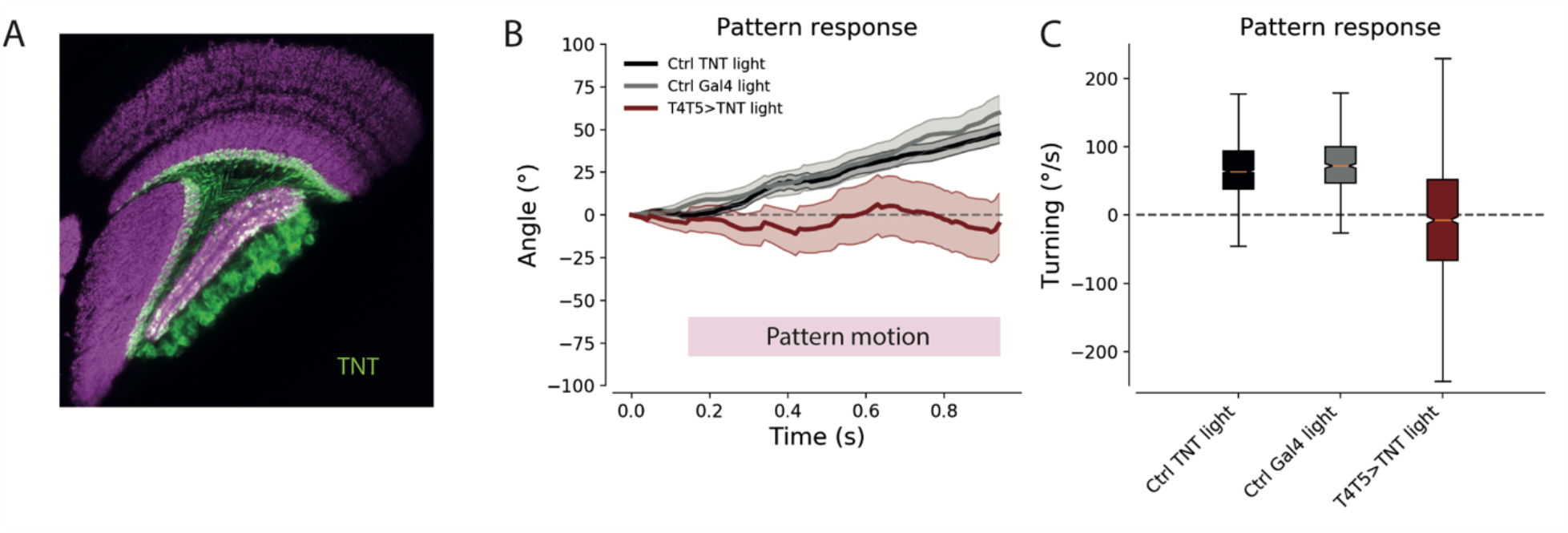
Response of T4/T5-block flies to pattern motion. (**A**) Confocal image of the split-Gal4-driver line (R42F06.AD; VTO43070.DBD) shown in a horizontal cross-section. Neurons are marked in green using an antibody against TNT, the neuropile is stained in purple by an antibody against the postsynaptic protein Dlg. (**B**) Turning response to pattern motion (mean ± s.e.m) of parental controls (TNT control n=419 trajectories - black, Gal4 control n=131 trajectories - grey) and motion blind flies (n=67 trajectories - dark red). Trajectories selected have minimum 150 ms of flight before pattern motion starts and at least 800 ms flight during pattern motion. (**C**) Average angular velocity of parental controls and motion blind flies during pattern motion (TNT control n=1622 trajectories –black, Gal4 control n=659 trajectories –grey, T4T5>TNT n=1333 trajectories – dark red).

We next analysed flight trajectories of control and T4T5>TNT flies in the presence of static patterns. Flight structure of T4T5>TNT flies appeared normal, albeit, an increased flight velocity was observed (Figure 3A) Intriguingly, inter-saccadic segment straightness of T4T5>TNT flies was reduced compared to controls. These results are similar to wild-type flies in the dark (compare Figure 3 with Figure 1E,F) demonstrating a role of motion vision for keeping flight trajectories straight.

**Figure 3.**
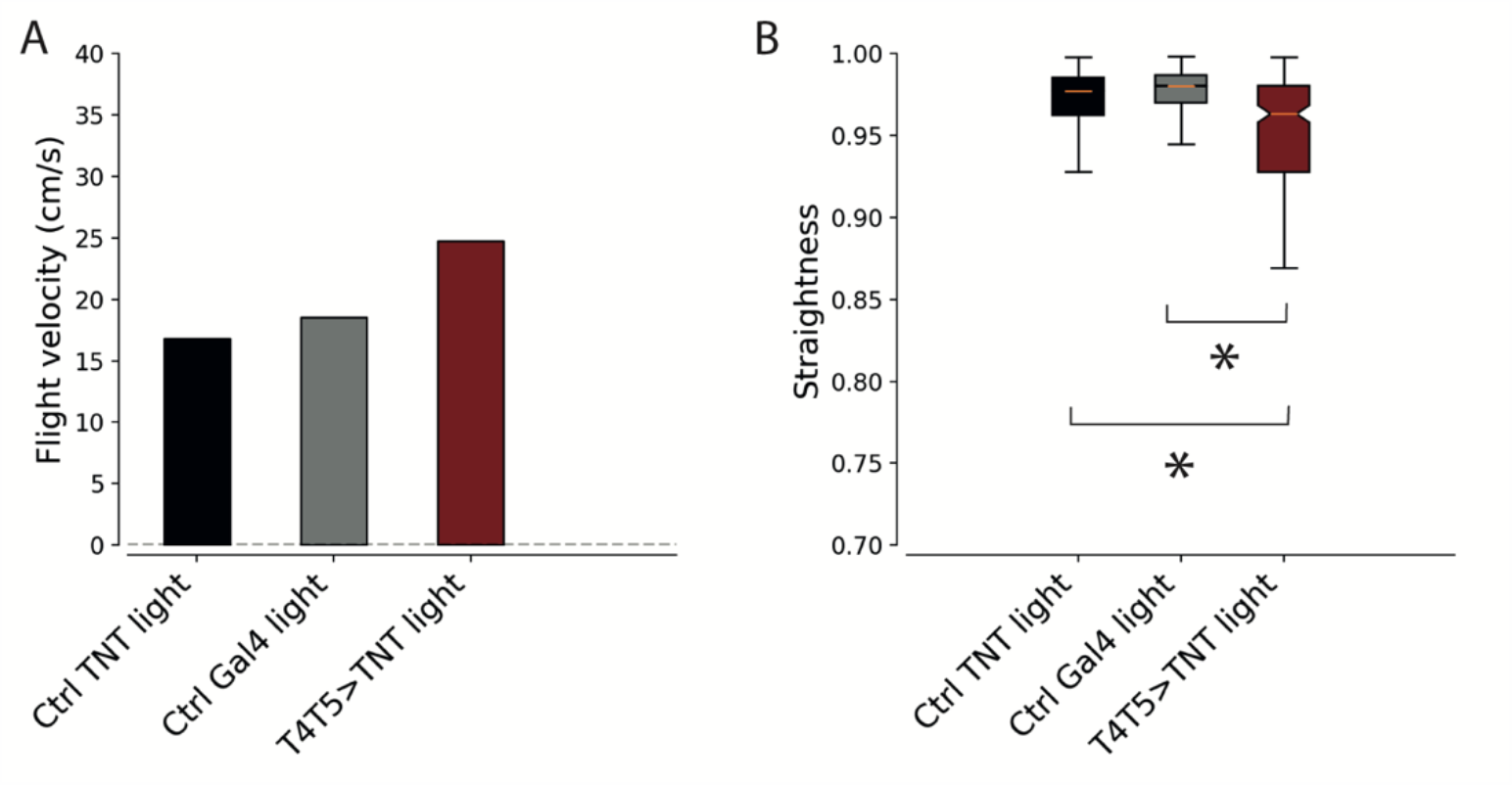
Free-flight behavior of motion-blind flies. (**A**) Flight velocity of parental controls and motion blind flies (s.e.m ≤ 0.043, TNT control n=2107 trajectories, Gal4 control n=2457 trajectories, T4T5>TNT n=512 trajectories). (**B**) Straightness of long intersaccadic flight segments (250 – 2000 ms) of parental controls and motion blind flies. Significance was calculated via a Kolgomorov/Smirnov test: p<0.0001 for each parental control vs T4T5>TNT flies (TNT control n=5842 segments, Gal4 control n=6612 segments, T4T5>TNT n=242 segments).

### Compensation of aerodynamic asymmetry

The above results suggest that the contribution of motion vision to keeping inter-saccadic flight straight is significant but subtle. However, inherent turning biases at the level of individuals (Souman et al., 2009) may be more pronounced under natural conditions than the average across the population of experimental flies tested in this study. Furthermore, asymmetric extrinsic forces such as air turbulences, prevalent in nature, were absent in our arena.

In order to test for the flies’ ability to compensate for a consistent turning bias, we clipped ∼25% of the tip of either the right or the left wing (Bender & Dickinson, 2006; Muijres et al., 2017). We processed data of left wing-clipped flies as if they were clipped on the right side, allowing us to combine data from both manipulations. We quantified trajectories from the following experimental groups: wildtype light and wildtype dark (Figure 4A-D), as well as TNT control light, Gal4 control light and T4T5>TNT light (Figure 4E-H). Visual inspection of individual trajectories (x/y coordinates) from wing-clipped wildtype flies in the light revealed a flight structure similar to intact controls (Figure 4A). However, wing-clipped wildtype flies in the dark behaved differently in that many flight trajectories exhibited a clockwise or counter-clockwise circular structure (Figure 4B). The same was true when comparing TNT controls (normal flight structure) with T4T5>TNT (curved trajectories) (Figure 4E, F).

**Figure 4.**
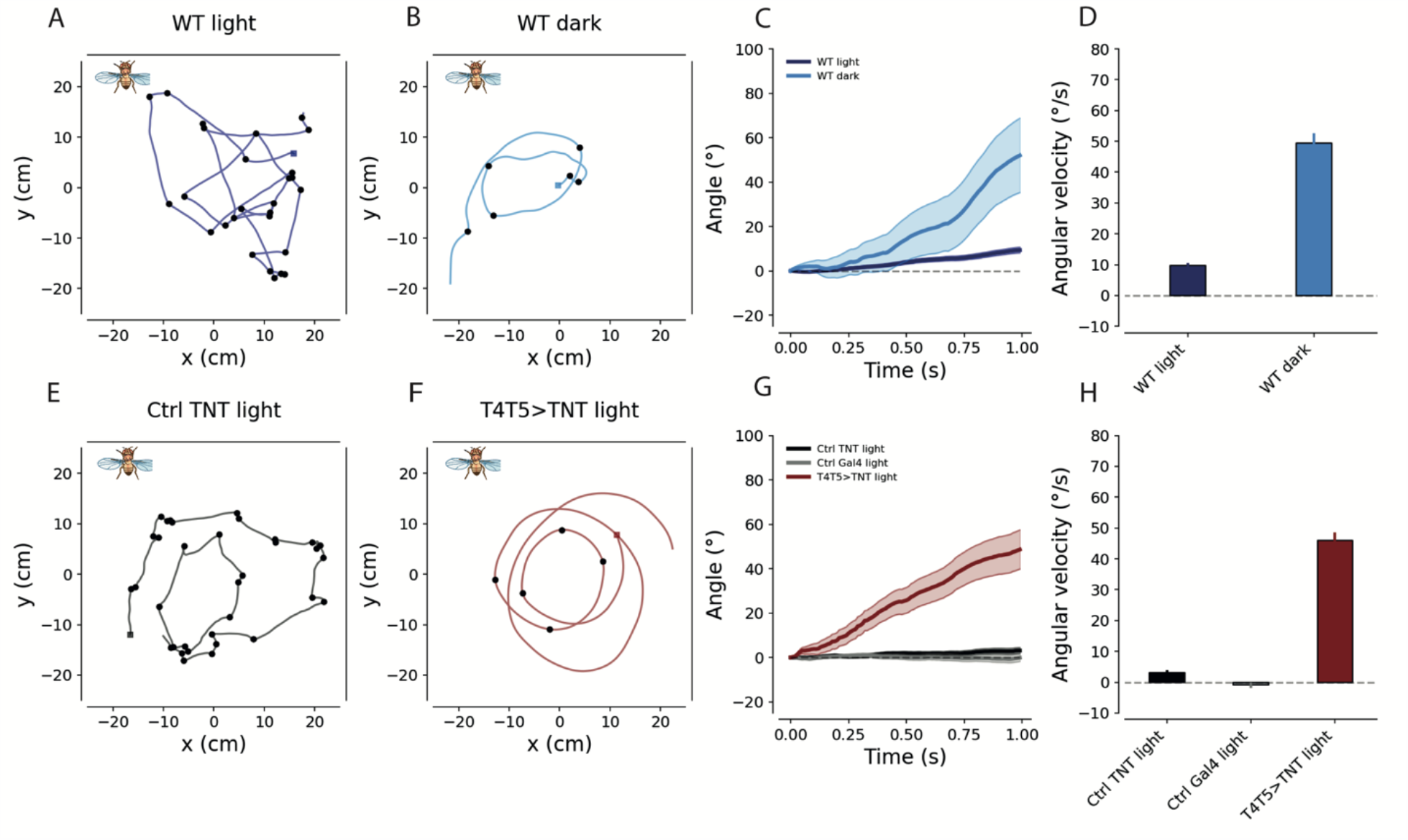
Compensation of aerodynamic asymmetry. (**A** and **B**) Example of a single flight trajectory of wild type fly with one clipped wing in the light (**A**, dark blue) and in the dark (**B**, light blue). (**C**) Mean flight angles (±s.e.m) over time of wild type flies with clipped wings (n=6773 segments light and n=205 segments dark). (**D**) Mean angular velocity (±s.e.m) of wild type flies with clipped wings (same dataset as in **C**). (**E** and **F**) Example of a single flight trajectory of a parental UAS control (**E**, black, grey) and a motion-blind fly (**F**, dark red) with clipped wings. (**G**) Mean flight angles of parental controls and motion blind flies with one clipped wing (TNT Control n= 4983, Gal4 Control n= 3893, T4T5>TNT n=734). (**H**) Mean angular velocity (±s.e.m) for parental controls and motion blind flies (same dataset as in **G**).

Calculating straightness indices revealed a strong reduction for wing-clipped wildtype dark and T4T5>TNT flies, compared to their respective controls (Figure S3). However, since saccade detection might be compromised by circling flight, we analysed trajectories independent of saccade detection in the following way: First, for each trajectory, we took the orientation over time, obtained from the angle of the vector defined by consecutive x-y-positions relative to the arena coordinates. We then subtracted the initial angle so that the orientation of each trajectory commenced at zero and computed the average across the first second of recording (Figure 4C, G). The average orientation of wildtype light, TNT control and Gal4 control flies revealed an almost perfect compensation, i.e. a small change in orientation over time. Both wildtype dark and T4T5>TNT flies, however, exhibited a pronounced average drift in the direction opposite to the wing-clipped side. Furthermore, taking the average across all recorded angular velocities resulted in values of 0 – 15 °/s for wildtype light, TNT control and Gal4 control flies. With 45 - 55 °/s, this parameter was much higher for wildtype dark and T4T5>TNT flies (Figure 4D, H).

To summarize, eliminating motion vision had the same effect as removing all visual input: flies lost their ability to compensate for an experimentally introduced turning bias. Hence, these results unequivocally demonstrate that motion vision contributes to course stabilization.

### Sensory cues complementary to optic flow

Animals have various additional sensory cues at their disposal to control heading. For instance, stable bearing can be aided by keeping conspicuous visual features stationary on the retina without requirement of explicit visual motion representation (Bahl et al., 2013). This involves the computation of an error angle to be minimized by appropriate turning reactions. In our arena, display monitor edges and buzzing devices may have provided flies with cues to determine their bearing. However, in nature, visual landmarks may not always be present as for instance in densely cluttered surrounds. Furthermore, using error angle for proportional control of heading is noise sensitive and prone to overshoot, as shown in bats (Bar et al., 2015). Optic flow in turn provides a signal akin to the derivative of an error angle (Bar et al., 2015). Under natural conditions, both position and motion vision system are likely used in a redundant fashion for robust steering.

In addition to vision, mechanosensory feedback from body appendages plays an important role to prevent accidental heading changes. In Diptera, for instance, club-shaped appendages modified from hind wings, the halteres, act as gyroscopes sensing body rotations (Nalbach, 1993; Dickinson 1999). Since they are tightly coupled to the wing motor system via afference and efference (Dickerson et al, 2019), they provide ultrafast feedback critical for stable flight. Visual motion in turn signals slower rotations (Sherman & Dickinson, 2004), complementing haltere feedback in a different angular velocity regime.

## Conclusion

It has long been recognized that motion vision is suitable to subserve various ethological functions. However, the significance of self-evoked visual motion signals for course control has been difficult to address. Here, by combining free-flight tracking with the specific removal of direction-selective neurons in flies, we firmly establish an important contribution of the motion vision system to course stabilization.

## Acknowledgements

We would like to thank T. Schilling for the characterization of T4/T5 Split-Gal4 driver lines, M. Sauter and J. Pujol-Marti for help with dissection and confocal imaging, S. Prech for technical assistance in building the free flight arena, C. Theile and R. Kutlesa for help with the tethered-walking experiment, W. Essbauer for fly work, B. Prud’homme, F. Schnorrer and N. Luis from University of Marseille, France for catching and providing the wild-caught strain ‘Luminy’, R. Strauss from University of Mainz, Germany for providing the UAS-TNT strain and W. Denk from the MPI of Neurobiology for carefully reading the manuscript. This work was supported by the Deutsche Forschungsgemeinschaft (SFB 870) and the Max Planck Society.

## Author contributions

A.L, A.B. and A.S.M conceived of the study. M-B.L., A.L, A.B. and A.S.M designed the experiments. A.L. set up the arena and wrote the tracking software. M-B.L. carried out all experiments. M-B.L. and A.S.M. analysed the data. A.S.M. wrote the manuscript with the help of all other authors.

## Declaration of Interests

The authors declare no competing interests.

## Material & Methods

### Flies

Flies were raised on standard fly food (cornmeal-agar) at 250C and 60% humidity, on a 12h light/12h dark cycle. The wild caught flies (‘Luminy’) were provided by Frank Schnorrer and collected by Benjamin Prud’homme. NorpA7 flies were obtained from Bloomington Stock Center (no. 5685). The split-Gal4 line used to drive expression in free flight experiments (w-/w-; R42F06.AD/Cyo; VT043070.DBD/Tm6b) was derived in our lab form AD and DBD domains found in Bloomington Stock Center (no. 70685 and no. 72763). This line was crossed to w+ cs; TNT-E/TNT-E; +/+ (from Roland Strauss) to create a fly line which blocks activity in all T4/T5 cells: w+/w-; R42F06.AD/TNT-E; VT043070.DBD/+. As controls, both the UAS and the split-Gal4 line were crossed with CantonS flies resulting in: w+/w-; R42F06.AD/+; VT043070.DBD/+ and w+ cs; TNT-E/+; +/+. The additional T4/T5 line used in experiments included in the supplemental data (w-/w-; UAS-Dcr2/Cyo; GMR39H12.Gal4/Tm6b) was derived from the driver line no. 50071 from Bloomington Stock Center.

Female flies younger than 48 hours were selected for free flight experiments using CO2 anesthesia. Wing ablation was performed using standard micro scissors (Fine Science Tools, art. no. 15001-08). The cut was executed chordwise to the wing using as landmark the spot where the Longitudinal Veins 1 and 2 connect at the distal part of the wing. The flies were moved in a new vial with standard food and placed back in the incubator for 24 hours to recover from CO2 exposure. They were then flipped into empty vials to induce starvation for 4 hours before being transferred to the experimental arena.

### Tethered walking

Assessment of optomotor response was done using a locomotion recorder previously described in Bahl et al. (2013). Briefly, a fly whose head, thorax and wings were fixed to a needle using near-ultraviolet bonding glue (Sinfony Opaque Dentin) and strong blue LED light (440 nm, dental curing light, New Woodpecker) was placed on an air-suspended sphere. The movements of the sphere were recorded via two optical tracking sensors. The experiments were performed at 340C, temperature achieved by a self-designed Peltier controlling system.

For each fly, the experiment consisted of 50 trials with stimuli in each trial being presented randomly. Gratings moving to either left or right were presented either for a short period of time (0.5 s) or long period of time (6 seconds) and either with a high or low contrast. Flies which walked continuously for at least 10 trials were selected and only the trials which had an average walking speed of higher than 0.25 cm/s were included in the analysis. Turning speed traces were determined by taking the average over trials and low-pass filtering the resulting trace (τ = 0.1 s in all experiments). All data analysis was performed in Python 3.7 using NumPy 1.15.1 and SciPy 1.1.0.

### Free-flight arena

The arena consisted of quadratic transparent enclosure (50×50×30 cm) of acrylic plastic (Evonik, Plexiglas XT). Below the bottom, through a diffuser, a custom-made array of high-intensity infrared LEDs (Roithner Laser Technik GmbH, H2A1-H830, peak at 830 nm far outside the activation spectrum of fly photoreceptors) provided strong background light to facilitate optical tracking by cameras with short exposure time and at high frequency (Point Grey Research, CM3-U3-13S2C-CS).

To encourage flight, small “buzzing devices” were placed in the four corners of the ceiling of the arena. Each device constituted a small Petri dish with yeast-supplemented fly food, i.e. emanating an attractive odour. Contact of flies with the food was prevented by a grid cover, to which a vibration motor (Pololu, #2265) was attached. Vibration went off every minute and evoked take-off and flight in flies sitting on the grid. This way, sufficient flight data was obtained even from flies in the dark, blind flies and motion-blind flies, which are otherwise reluctant to fly.

Static or moving visual patterns were displayed on four monitors (ASUS, VG248QE) placed on the sides of the enclosure at 120 Hz. A regular checkerboard pattern of 5×5cm in checker size was displayed as a static visual stimulus for the majority of the experiments. For the experiment shown in Figure 2, the same pattern would move at 40cm/s to the left for 10 seconds, it would remain static for another 10 seconds and then would move to the right for 10 seconds, followed by another 10 s of static display.

### Free-flight tracking

#### Cameras

Our multi-camera set-up consisted of 5 mounted units (FLIR Inc., CM3-U3-13Y3M-CS) observing overlapping volumes of the arena (see illustration in Fig. 1 B). We used standard machine vision lenses with a focal length of approximately 6 mm (Thorlabs Inc., MVL6WA). Cameras were connected to a single tracking computer via USB3. To guarantee accurate synchronization across frame captures, image acquisition for all units was triggered by a single external TTL pulse generator that ran at 100 Hz. To prevent leakage of the visual stimulus into tracking images, we equipped all cameras with near-IR longpass filters (Thorlabs Inc., FGL780M) that separated the displays’ spectrum from near-IR background lighting.

#### Camera calibration

We calibrated intrinsic and extrinsic camera parameters with a single-step method (Li et al, 2013) that estimates the relevant matrices of all units in a multi-camera set-up from overlapping presentations of a printed random calibration pattern. The underlying camera model was a standard pinhole camera (with radial distortion). On average, we were able to achieve a reprojection error below 1 px based on 100-200 synchronized multi-camera snapshots of the calibration pattern. We used the algorithm as implemented in the provided MATLAB toolbox (https://sites.google.com/site/prclibo/toolbox). Calibration was performed periodically throughout the experimental phase to safeguard against shifts in camera position that could affect triangulation.

#### Detection

We processed incoming images from 5 cameras running at 100 Hz. Images were acquired as 640 x 512 px single-channel matrices. We estimated the static background by accumulating incoming images with a weight of 0.001 on each time step and subtracted this background estimate from new frames to isolate moving targets. Finally, we thresholded the resulting image at a minimum value of 5 (out of 255) to further suppress photon noise. With each camera image, we applied a standard blob detection algorithm from OpenCV (“cv2.findCountours”) to detect contiguous 2D targets. The position of a target was then defined as the weighted center of the blob. Coordinates of these targets were combined in the triangulation process described below.

#### Reconstruction

Unlike previous studies (Straw et al., 2011), we separated the tracking step into per-frame triangulation and subsequent association to generate trajectories for identified individuals. Triangulation was accomplished through application of the Hungarian algorithm (Ardekani et al., 2013).

Briefly, for each frame our 2D tracking algorithm yields multiple x-y detections which need to be associated across cameras to allow correct reconstruction of 3D positions. If only a single fly moves inside the arena, this operation is trivial; we find a single 2D detection per camera and all detections emanate from the same target. However, when multiple flies move simultaneously we need to correctly assign 2D detections to 3D targets. We always use standard singular value decomposition to estimate the optimal 3D position from a set of 2D noisy observations (Hartley & Zissermann, 2003). The Hungarian algorithm then efficiently calculates a minimum-cost assignment where cost is defined as the reprojection error after assigning particular 2D points to particular 3D targets. We implemented the algorithm in Python 2.7 and Numba, relying on OpenCV or PyMVG (https://github.com/strawlab/pymvg) for various projection operations.

#### Filtering and association

The method outlined above provides a number of 3D targets per frame. Per-individual analysis requires an association step where these points are aggregated into defined trajectories of single flies. The tracking algorithm treats targets as a collection of linear Kalman filters. Observations are 3D positions. The underlying state consists of six parameters; instantaneous 3D position as well as three velocities in all three directions. All filters are based on a constant-velocity process where maneuvering is modeled as noisy deviations from this constant velocity. The process matrix simply advances the current position by estimated velocity times the frame length (10 ms). We assume the following standard deviations for the different components: 2 cm for the measurement noise as well as 1 cm and 50 cm s-1 for the position and velocity components of the process noise, respectively. Standard deviations for the state covariance matrix are initialized as 10 cm for x-y-z position and 100 cm s-1 for all velocities. We did not tune these parameters extensively as they had little effect on tracking quality.

On each time step, we predict the position of each target and use the Hungarian algorithm to assign novel 3D observations to the set of existing filters (based on aggregated distance cost of the assignment). Observations can only be assigned to a filter if the distance is below 1 cm. Any observation that cannot be matched to an existing filter spawns a new target. If a filter does not receive a fresh observation for 20 time steps, the instance is terminated. A trajectory is then simply the filtered position estimate of a single Kalman instance from spawning to termination.

No additional post-processing was applied to disambiguate crossing paths of flies as we found these events to be rare in practice. Reconstruction of 3D points and tracking were computed offline. We used Python 2.7 and the filterpy package (https://filterpy.readthedocs.io/en/latest/) to implement these routines.

### Free-flight data analysis

#### Trajectory selection and feature extraction

All analysis was carried out in Python 3.7, using the following libraries (among others): NumPy 1.17.2, Pandas 0.25.1, SciPy 1.3.1. Movement trajectories (walking and flight) were loaded and those below a minimal length (1s) discarded. Values for x, y, and z were smoothed by convolving them separately with a block filter of size 9.

To obtain an initial selection of flight trajectories, only segments of length > 1s within a certain z range (1.5 cm above floor and 1.0 cm below ceiling) were included and labelled with a new identifier. From positions over time, the following features were extracted: x-y angle (°), x-y angular velocity (°/s), x-y-z flight velocity (cm/s) and x-y (horizontal) flight velocity (cm/s). At this point, manual inspection still revealed a fraction of walking trajectories based on low movement velocity and little x-y displacement over time. Hence, trajectories with a mean flight velocity below 3.0 cm/s or the sum of x and y standard deviation below 2 cm were discarded. Saccade detection. Saccade detection was carried out based on angular velocity, obtained by differentiating the angle of consecutive x/y positions relative to the arena coordinates. For each flight trajectory, angular velocity was convolved with a gaussian kernel of the approximate shape of a saccade (σ = 40 ms). Saccade time points were then identified by peak values above a threshold of 300 °/s. This procedure was done separately for left- and rightward saccades, taking the respective sign into account.

#### Straightness

For each intersaccadic segment, distance and path length between the two endpoints was computed. To obtain the straightness index, distance was divided by path length. Statistical significance was established by computing the Kolmogorov-Smirnov statistic on 2 samples using scipy.stats.ks_2samp in Python.

### Immunohistochemistry

Primary antibodies used: mouse anti-Bruchpilot (1:20, Developmental Studies Hybridoma Bank, AB2314866), rabbit anti-Tetanus Toxin (1:5000, SSI Antibodies 65873 POL 016). Secondary antibodies used: ATTO 647N goat anti-mouse (1:400, Rockland 610-156-040), Alexa Fluor 568-conjugated goat anti-rabbit (1:400, Life Technologies A-11011).

Brains were dissected in cold PBS and fixed in 4% paraformaldehyde (0.1% Triton X-100) for 25 minutes at room temperature. They were then washed 3 times with PBST (PBS containing 0.3% Triton X-100) and blocked with Normal Goat Serum (10% NGS in PBST) for 1 hour. Brains were incubated at 40C for 48 hours with primary antibodies diluted in the NGS solution previously described. They were washed 3 times (for 1-2 hours each) with PBST and then incubated at 40C for 48 hours with secondary antibody diluted in the NGS solution. Brains were then washed three times in PBST before mounting in SlowFade Gold Antifade Mountant (Thermo Fisher Scientific).

---

**Figure S1.**
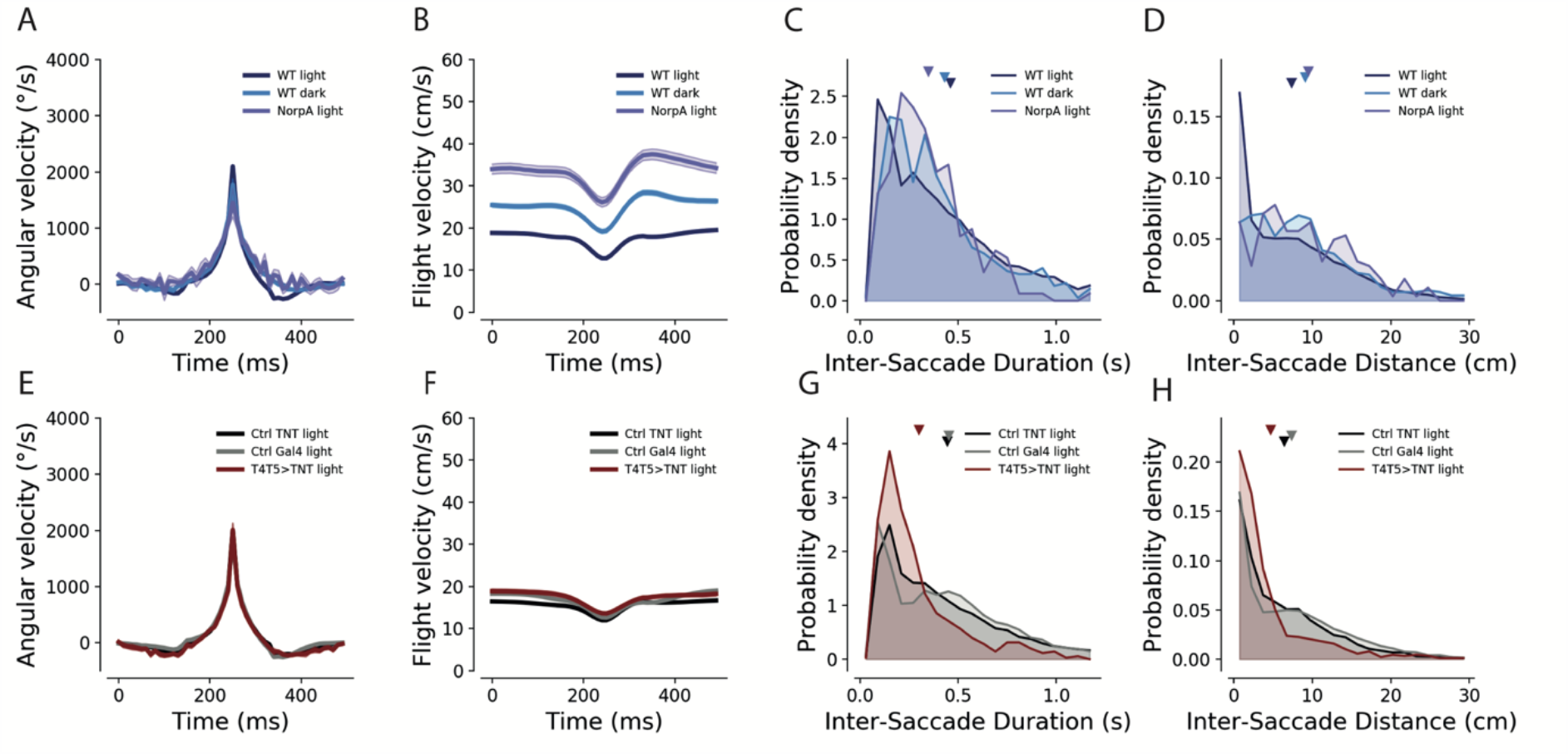
Saccade characteristics across genotypes and stimulus conditions. (A and E) Angular velocity profile during a saccade. (B and F) Flight velocity during a saccade. (C and G) Probability density of the duration between saccades. (D and H) Probability density of distance between saccades.

**Figure S2.**
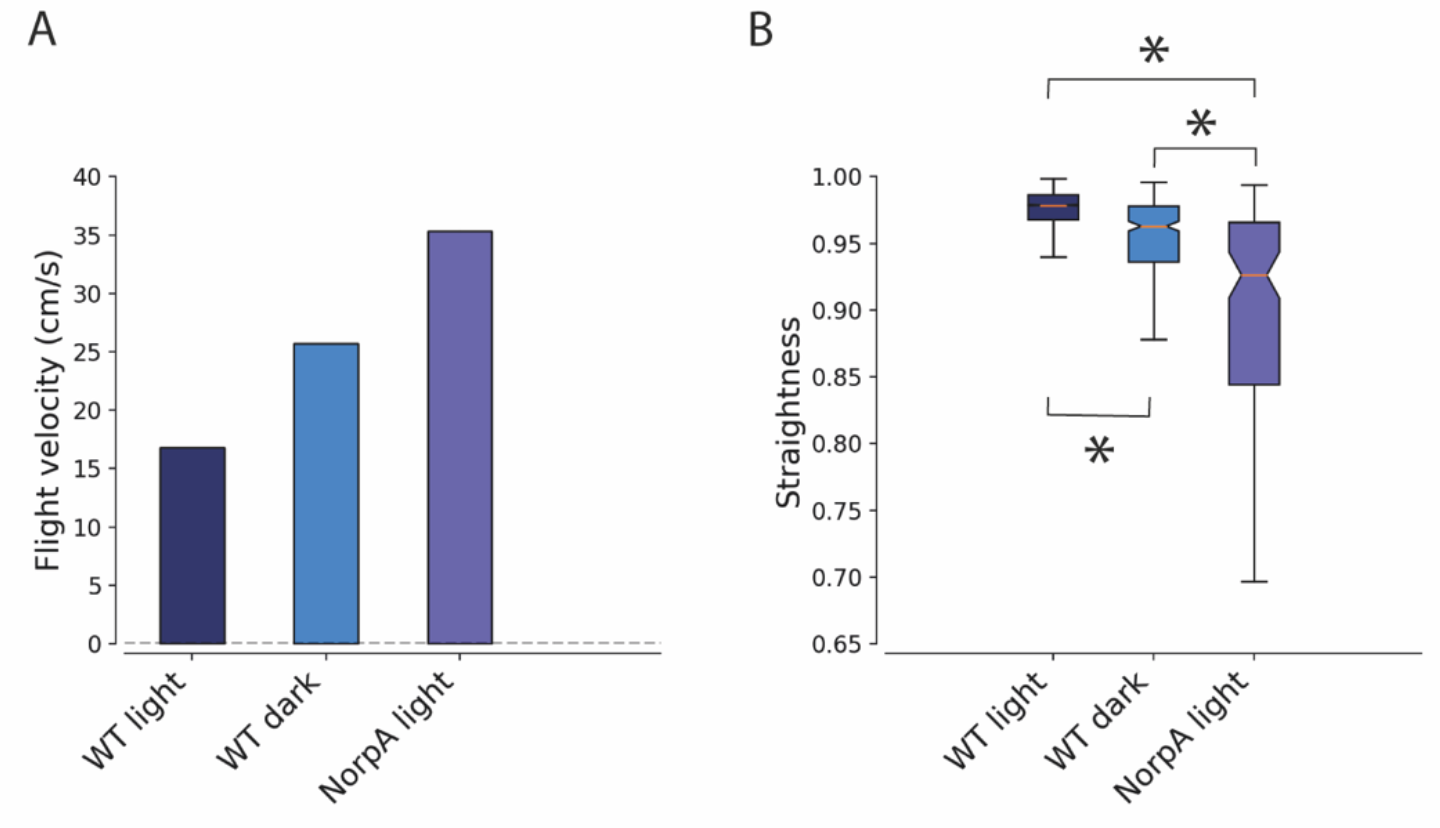
Free-flight behavior of wildtype flies in the light and in the dark as well as of blind flies. (A) Average flight velocity (s.e.m ≤ 0.045) of flies in the light, in the dark and blind flies (n=1988 trajectories light, n=310 trajectories dark, n=100 trajectories blind flies). (B) Straightness of long inter-saccade flight segments (250-2000 ms). Significant differences based on a Kolgomorov/Smirnov test (p<0.0001, n=4609 segments light, n= 302 segments dark, n=100 segments blind).

**Figure S3.**
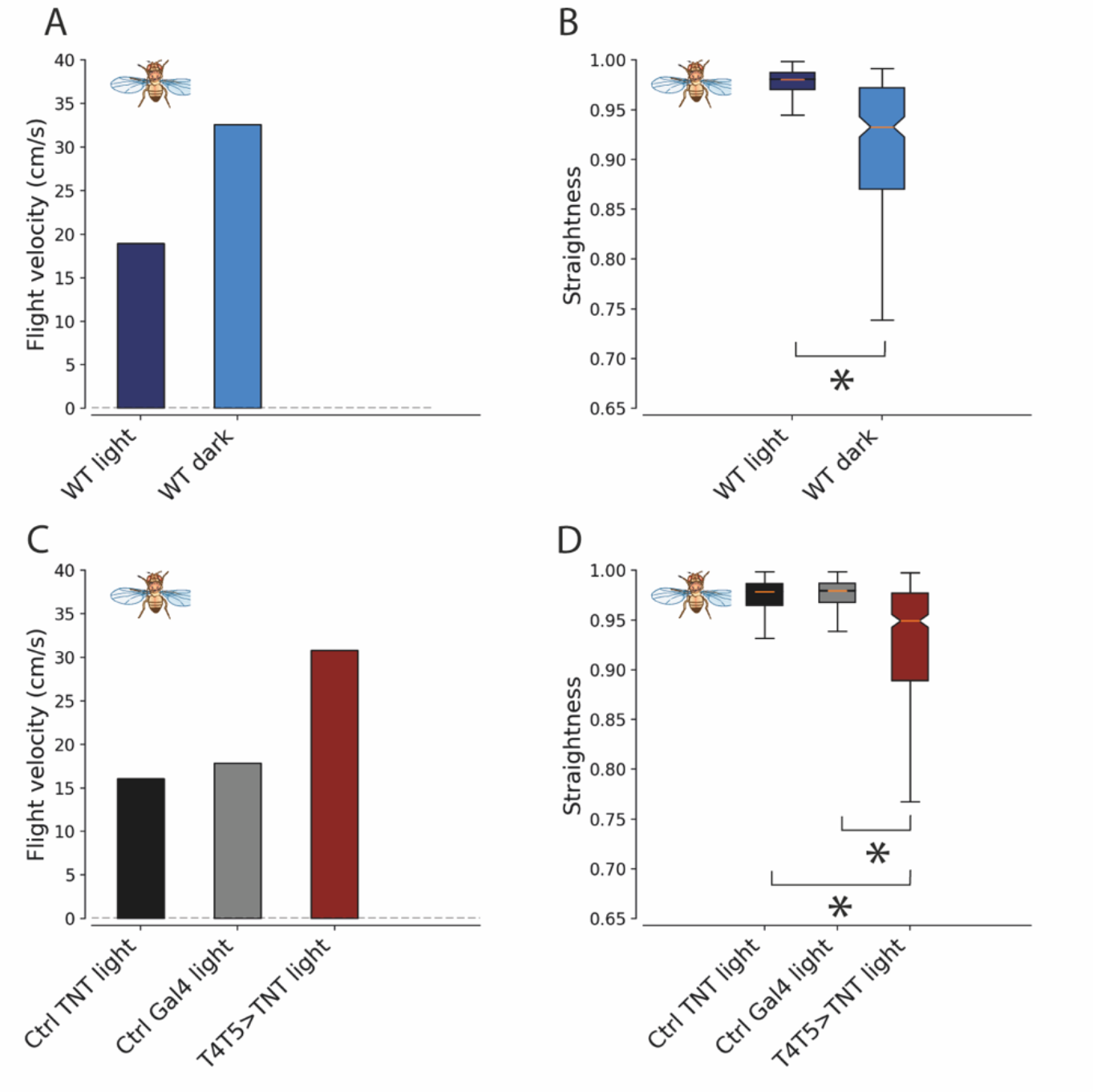
Free-flight behavior of wing-clipped flies. (A) Average flight velocity (s.e.m ≤ 0.077) of flies in the light and in the dark (n=6773 trajectories light and n= 205 trajectories dark). (B) Straightness of inter-saccade flight segments (250 – 2000 ms) of wildtype flies. Significant differences based on Kolgomorov/Smirnov test (n=7607 inter-saccade flight segments light and n=241 segments dark p<0.0001). (C) Average flight velocity (s.e.m ≤ 0.048) of motion-blind flies and parental controls (TNT Control n=4983 trajectories, Gal4 Control n=3893 trajectories, T4T5>TNT n=734 trajectories). (D) Straightness of inter-saccade flight segments (250 – 2000 ms) of parental control and T4T5 > TNT flies. Significant differences are based on Kolgomorov/Smirnov test (p<0.0001, TNT Control n= 6818 inter-saccade flight segments, Gal4 Control n= 7162 segments, T4T5>TNT n=418 segments).

**Figure S4.**
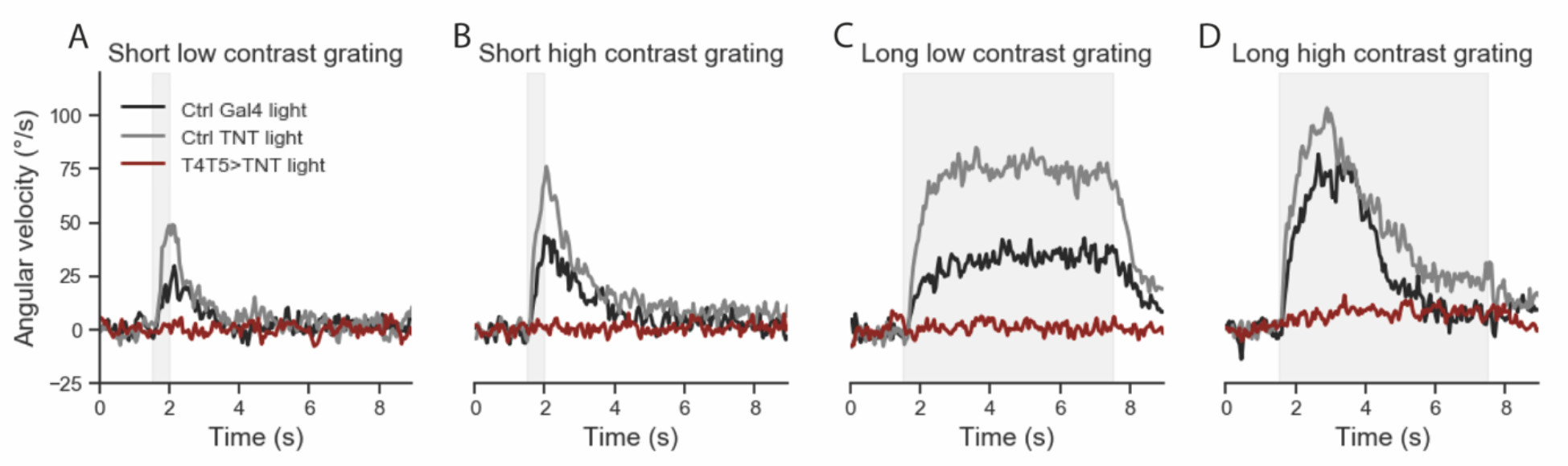
Angular velocity of tethered-walking flies during presentation of moving grating stimuli with different contrasts and duration. (A) Angular velocity response to short low contrast grating. (B) Angular velocity response to short high contrast grating. (C) Angular velocity response to long low contrast grating. (D) Angular velocity response to long high contrast grating. n=7 motion blind flies, n=9 for TNT Control, n=4 Gal4 Control flies

## Notes

### Competing Interest Statement

The authors have declared no competing interest.

